# Whole-brain connectivity during encoding: age-related differences and associations with cognitive and brain structural decline

**DOI:** 10.1101/2021.08.10.455779

**Authors:** Elettra Capogna, Markus H. Sneve, Liisa Raud, Line Folvik, Hedda Ness, Kristine B. Walhovd, Anders M. Fjell, Didac Vidal-Piñeiro

**Author notes:** **Corresponding author: Elettra Capogna**, Department of Psychology, Pb. 1094, Blindern, Oslo, Norway, 0317.

## Abstract

There is a limited understanding of age differences in functional connectivity during memory encoding. In the present study, a sample of cognitively healthy adult participants (n=488), a subsample of whom had longitudinal cognitive and brain structural data spanning 8 years back, underwent fMRI while performing an associative memory encoding task. We investigated 1) age changes in whole-brain connectivity during memory encoding; whether 2) encoding connectivity patterns overlap with the activity signatures of specific cognitive processes and whether 3) connectivity changes associated with memory encoding related to longitudinal brain structural and cognitive changes. Age was associated with decreased intranetwork connectivity and increased connectivity during encoding. Task-connectivity between mediotemporal and posterior parietal regions – which overlapped with areas involved in mental imagery – was related to better memory performance only in older age. The connectivity patterns supporting memory performance in older age reflected preservation of thickness of the medial temporal cortex. These investigations collectively indicate that functional patterns of connectivity should be interpreted in accordance with a maintenance rather than a compensation account.

## 1. Introduction

Episodic memory declines with age (Rönnlund et al., 2005), although there is substantial inter-individual variability in the trajectories (Nyberg et al., 2012). Variability in memory function is affected by changes in the structural and functional architecture of the brain. Hence, how brain regions communicate during memory tasks may be a key factor in explaining age-related changes in memory performance as well as inter-individual variation of performance in older age. Assessing brain functional connectivity changes in task contexts provides a window for studying age-related brain changes in response to specific cognitive demands (Campbell and Schacter, 2017). In the present study, we used task-related functional magnetic resonance imaging (fMRI) in a large adult lifespan sample to investigate whether 1) age is associated with changes in task-connectivity during encoding, specifically with decreased intranetwork and increased internetwork connectivity; 2) encoding connectivity patterns overlap with the activity signatures of specific cognitive processes; 3) connectivity changes associated with memory encoding performance relate to longitudinal brain structural and cognitive changes.

Most previous studies on functional connectivity changes in aging have employed seed-based task-functional connectivity (Grady et al., 2003; Oh and Jagust, 2013), or resting-state fMRI (rs-fMRI) (Fjell et al., 2015). Seed-based task-connectivity studies have repeatedly found higher age to be related to greater connectivity between medial temporal lobe, most notably the hippocampus, and prefrontal areas, during encoding (Grady et al., 2003; Oh and Jagust, 2013). Research using rs-fMRI has found lower intranetwork connectivity with higher age - especially within the default-mode regions - and increased connectivity between networks such as the dorsal attention and the default-mode network (Sala-Llonch et al., 2015; Vidal-Piñeiro et al., 2014). However, encoding connectivity exhibits a substantially different pattern from that observed during rest (Keerativittayayut et al., 2018). Therefore, it is crucial to understand how whole-brain task-connectivity during memory encoding contributes to successful recollection. Encoding-based connectivity has been characterized by increased communication between distant areas, such as higher integration of default mode, salience, and subcortical networks with the other subnetworks (Keerativittayayut et al., 2018). Furthermore, flexible nodes – nodes that change networks membership during different episodic memory task phases – appear to be relevant for memory performance as degree of observed reorganization between states partially predicts retrieval success (Schedlbauer and Ekstrom, 2019). A small number of studies have tested age-related differences in whole-brain connectivity during memory encoding (Grady et al., 2016; Matthäus et al., 2012; Wang et al., 2010). Wang and colleagues (2010) found higher age to be associated with lower long-range functional connections of frontal regions with the rest of the brain during encoding. Matthaus et al. (2012) observed age-related increases in the density and size of the networks together with reduced efficiency of information processing during encoding.

Age-related functional changes may accompany brain structure decline (Nyberg et al., 2012). For instance, in accordance with the brain maintenance framework, older adults showing longitudinal changes in prefrontal activity – and in further areas beyond task-specific regions – exhibited greater memory and hippocampal volume decline (Persson et al., 2006; Pudas et al., 2018). Alternatively, age-differences in connectivity may reflect an attempt to compensate for neural breakdown (Cabeza et al., 2018). For instance, one study found that higher connectivity between the prefrontal cortex with the rest of the brain was related to sustained memory performance uniquely in older adults (Deng et al., 2021), supporting a compensatory account for the age-related connectivity changes. In general, coupling age-related differences in function with cross-sectional performance is however not without problems, as compensatory responses can lay anywhere along a continuum from (partial) failure to success (Grady, 2012). Hence, for a better and complete understanding, functional differences need to be associated with brain and cognitive changes assessed over time.

Here, we investigated age-related differences in functional connectivity during an associative encoding task using a correlational psychophysiological interaction (cPPI) approach (Fornito et al., 2012) and a sample encompassing the entire adult age range. We assessed connectivity changes during encoding associated with age, memory performance, and the interaction between age and memory. Specifically, we focused on whether age was associated with decreased intranetwork and increased internetwork connectivity during the task, as typically shown in whole-brain resting-state (Geerligs et al., 2015) and in ROI-based task-connectivity studies (Grady et al., 2016; Spreng et al., 2016). Moreover, by comparing connectivity maps with meta-analytic activity maps, we investigated whether encoding connectivity patterns overlapped with the activity signatures of specific cognitive processes. Finally, among older adults, we tested whether the connectivity changes associated with successful memory encoding were related to longitudinal structural and cognitive changes. This allowed us to test whether these functional patterns of connectivity should be interpreted in accordance with the maintenance or the compensation accounts.

## 2. Material and methods

### 2.1 Participants

A total of 488 participants (336 females, mean age = 41.65 years, SD = 17.20, age range = 18-81) were included in the final sample. All participants completed the fMRI tasks and were screened through health and neuropsychological assessments. Participants were required to have no history of neurological or psychiatric disorders, chronic illness, be right-handed, and not to use medicines known to affect nervous system functioning. Participants were further excluded based on the following neuropsychological criteria: score <26 on the Mini-Mental State Examination (MMSE) (Folstein et al., 1975), score <85 on the WASI II (Wechsler, 1999), and a T-score of ≤30 on the California Verbal Learning Test II—Alternative Version (CVLT II) (Delis et al., 2000) immediate delay and long delay. All participants gave written informed consent, and the study was approved by the Regional Ethical Committee of South Norway and conducted in accordance with the Helsinki declaration. Retrospective longitudinal data were available for a subsample of older participants (age > 50 years), spanning up to 10 years back as follows: neuropsychological testing for 151 participants (n = 51, 6, and 94 with 1, 2, and ≥ 3 observations, respectively), and brain structural scans for 88 participants (n= 2, 5, and 81 with 1, 2, and ≥ 3 observations, respectively). Note that a small subsample of participants had retrospective data acquired with a different scanner. See **Supplementary Table 1** for more information.

### 2.2 Experimental design and behavioral analysis

The experiment included an incidental encoding task and a memory test after approximately 90 minutes, both in the scanner. In this study, we only analyzed encoding fMRI data. The experimental design has been thoroughly described elsewhere (Sneve et al., 2015; Vidal-Piñeiro et al., 2019). See **Fig. 1** for a visual description of the experiment. In brief, the encoding and retrieval tasks consisted of 2 and 4 runs, respectively, that included 50 trials each. The stimulus material comprised 300 black and white line drawings of everyday items. A central fixation cross was shown during the baseline recording at the beginning, the middle, and the end of each run for 11 seconds. In the encoding phase, the trial started with a voice asking the participants either “Can you eat it?” or “Can you lift it?”. Each question was asked 25 times in each run in a pseudorandomized order. One second after the question, an item appeared on the screen for 2 seconds, asking the participant to answer “Yes” or “No”, before being replaced by a fixation cross that remained throughout the intertrial interval (between 1 and 7 seconds, exponential distribution; duration = 2.98 [2.49] seconds). In the retrieval phase, the trial started with Question 1: “Have you seen this item before?”. The item appeared on the screen for 2 seconds, and the participant had to press “Yes” (old item), or “No” (new item). In each run, 25 old items and 25 new items were presented in a pseudorandomized order. If the participant responded “No”, the trial ended. If the participant responded “Yes”, the trial proceeded to Question 2: “Can you remember what you were supposed to do with the item?”. Again, if the participant responded “No”, the trial ended, if “Yes” the trial continued with Question 3: “Were you supposed to eat it or lift it?”. The participant was forced to choose between the two actions associated with the item at encoding. For behavioral analysis, the classification of responses to old items was: (1) source memory (“Yes” response to Question 1 and Question 2, and correct answer to Question 3), (2) item memory (“Yes” response to Question 1 and either “No” to Question 2 or incorrect answer to Question 3), (3) miss (incorrect answer to Question 1). New items were classified either as (4) correct rejections or (5) false alarms. Memory performance in the task was calculated as the proportion of source memory minus incorrect judgments to Question 3, tentatively controlling for false memories and guessing behavior (Vidal-Piñeiro et al., 2019). The relationship between age and relevant behavioral and neuropsychological metrics was tested with generalized additive models (GAMs), controlling for sex.

**Fig.1.**
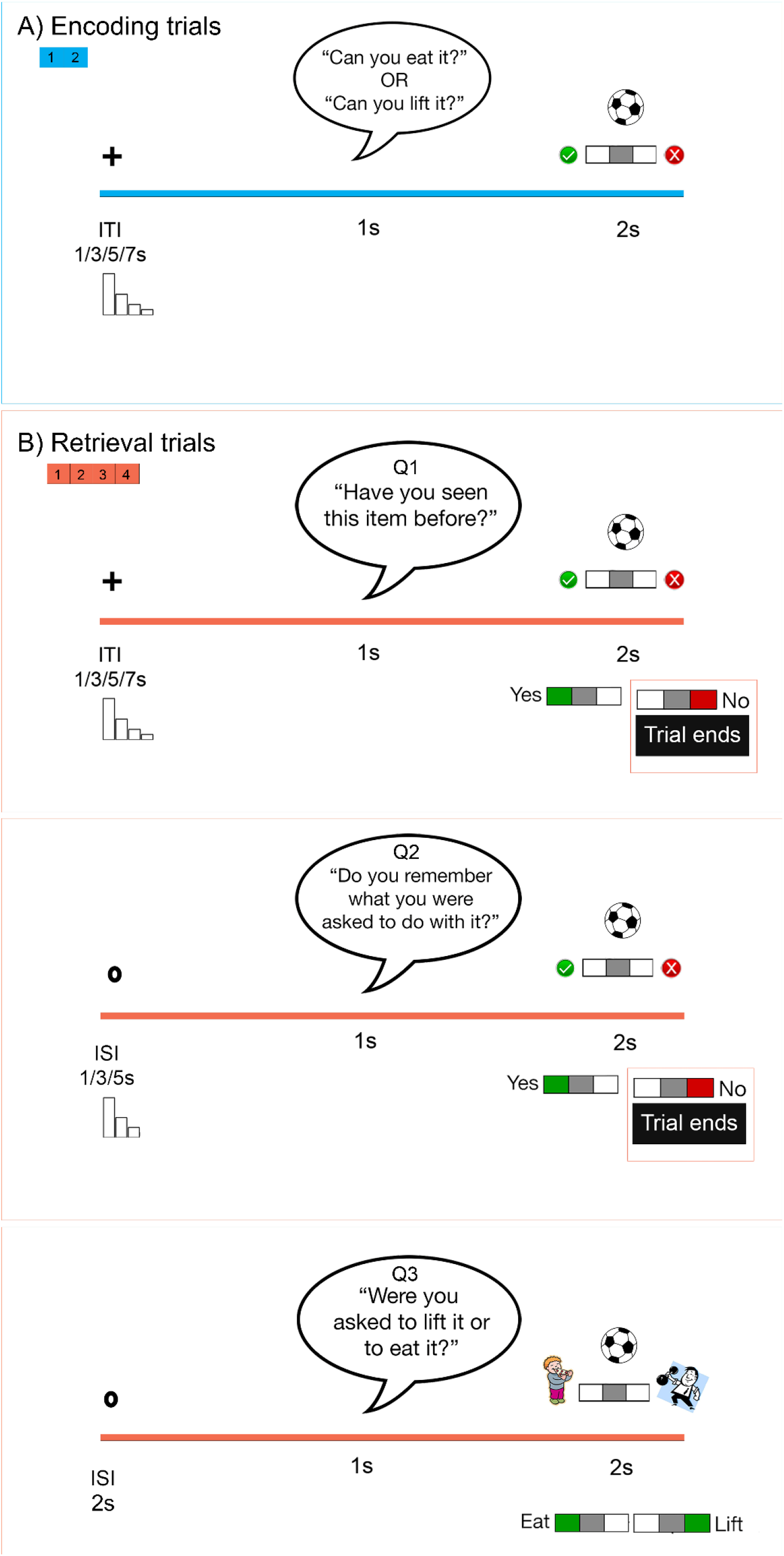
Experimental paradigm. **A** One trial of the encoding task. The green V and the red X were present on the screen to indicate which button indicate Yes and No. **B** One trial of the retrieval task. Test Questions 1 and 2 required a Yes/No response, whereas Question 3 required a choice between the two actions. The trial ended if the participant responded No to either one of the two first questions. Response cues (V, X, eating, lifting) were also present on the screen. ITI intertrial interval, ISI interstimulus interval. Adapted from Vidal-Piñeiro and colleagues (2017).

### 2.3 MRI acquisition

Imaging data were collected using a 20-channel Siemens head-neck coil on a 3T MRI (Siemens Skyra Scanner, Siemens Medical Solutions, Germany) at Rikshospitalet, Oslo University Hospital. The functional imaging parameters were equivalent across all fMRI runs: 43 transversally oriented slices (no gap) were measured using a BOLD-sensitive T2*-weighted EPI sequence (TR = 2390 ms, TE = 30 ms, flip angle = 90°, voxel size = 3 x 3 x 3 mm^3^, FOV = 224 × 224 mm^2^, interleaved acquisition; generalized autocalibrating partially parallel acquisitions acceleration factor [GRAPPA] = 2). Each encoding run produced 134 volumes. At the start of each fMRI run, 3 dummy volumes were collected to avoid T1 saturation effects. Anatomical T1-weighted (T1w) magnetization-prepared rapid gradient echo (MPRAGE) images consisting of 176 sagittally oriented slices were obtained using a turbo field echo pulse sequence (TR = 2300 ms, TE = 2.98 ms, TI = 850 ms, flip angle = 8°, voxel size = 1 × 1 × 1 mm^3^, FOV = 256 × 256 mm^2^) were also acquired. Furthermore, a standard double-echo gradient-echo field map sequence was acquired for distortion correction of the echo planar images. Visual stimuli were presented in the scanner environment with an NNL 32-inch LCD monitor while participants responded using the ResponseGrip device (both NordicNeuroLab, Norway). Auditory stimuli were presented to the participants’ headphones through the scanner intercom. The structural T1w data used in the longitudinal analysis were collected using a 12-channel head coil on a 1.5 T Siemens Avanto scanner (Siemens Medical Solutions, Germany) at Rikshospitalet, Oslo University Hospital. The pulse sequence acquired consisted of two repeated 160-slice sagittal T1-weighted MPRAGE sequences (TR = 2400 ms, TE = 3.61 ms, TI = 1000 ms, flip angle = 8°, voxel size =1.25 × 1.25 × 1.20 mm, FOV = 240 mm). The raw images were automatically corrected for spatial distortion due to gradient nonlinearity (Jovicich et al., 2006) and field inhomogeneity (Sled et al., 1998), averaged, and resampled to isotropic 1 mm voxels.

### 2.4 MRI preprocessing

#### 2.4.1 fMRI preprocessing

Data were organized and named according to the Brain Imaging Dataset Specification standard (*BIDS*) and preprocessed using a *fMRIPep* preprocessing pipeline (Esteban et al., 2019) (v. 1.2.5) a “Nipype” based tool (Gorgolewski et al., 2018) (v. 1.1.6).

The T1w image was corrected for intensity non-uniformity (INU) using *N4BiasFieldCorrection* (Tustison et al., 2010) (ANTs v. 2.2.0), and used as T1w-reference throughout the workflow. The T1w-reference was then skull-stripped using *antsBrainExtraction.sh* (ANTs v. 2.2.0), using OASIS as target template. Brain surfaces were reconstructed using recon-all (FreeSurfer v. 6.0.1) (Dale et al., 1999), and the brain mask estimated previously was refined with a custom variation of the method to reconcile ANTs-derived and FreeSurfer-derived segmentation of the cortical gray matter (GM) of *Mindboggle* (Klein et al., 2017). Spatial normalization to the ICBM152 Nonlinear Asymmetrical template version 2009c (Fonov et al., 2009) was performed through nonlinear registration with *antsRegistration* (Avants et al., 2008), using brain-extracted versions of both T1w volume and template. Brain tissue segmentation of cerebrospinal fluid, white-matter and grey matter was performed on the brain-extracted T1w using FAST (FSL v. 5.0.9) (Zhang et al., 2001).

For each BOLD run, the following preprocessing was performed: first, a reference volume and its skull-stripped version were generated using a custom methodology of fMRIPrep. A deformation field to correct for susceptibility distortions was estimated based on a field map that was co-registered to the BOLD reference, using a custom workflow of fMRIPrep derived from D. Greve’s *epidewarp.fsl* script and further improvements of HCP Pipelines (Glasser et al., 2013). Based on the estimated susceptibility distortion, an unwarped BOLD reference was calculated for a more accurate co-registration with the anatomical reference. The BOLD reference was then co-registered to the T1w reference using *bbregister* (FreeSurfer). Co-registration was configured with six degrees of freedom. Head-motion parameters with respect to the BOLD reference (transformation matrices, and six corresponding rotation and translation parameters) were estimated before any spatiotemporal filtering using *mcflirt* (FSL v. 5.0.9) (Jenkinson et al., 2002). BOLD runs were slice-time corrected using *3dTshift from AFNI v. 20160207* (Cox and Hyde, 1997). The BOLD time-series (including slice-timing correction when applied) were resampled onto their original, native space by applying a single, composite transform to correct for head-motion and susceptibility distortions. These resampled BOLD time-series will be referred to as preprocessed BOLD in original space, or just preprocessed BOLD. Several confounding time-series were calculated based on the preprocessed BOLD: framewise displacement (FD) was calculated for each functional run, using Nipype’s implementation (following the definitions by Power and colleagues (2014)). Additionally, a set of physiological regressors were extracted to allow for component-based noise correction (*CompCor*) (Behzadi et al., 2007). Principal components were estimated after high-pass filtering the preprocessed BOLD time-series (using a discrete cosine filter with 128s cut-off). A subcortical mask was obtained by heavily eroding the brain mask to ensure it would not include cortical grey matter regions. Six anatomical CompCor (aCompCor) components were then calculated within the intersection of the aforementioned mask and the union of cerebrospinal fluid and white matter masks calculated in T1w space, after their projection to the native space of each functional run (using the inverse BOLD-to-T1w transformation). The head-motion estimates calculated in the correction step were also placed within the corresponding confounds file. All resamplings were performed with a single interpolation step by composing all the pertinent transformations (i.e., head-motion transform matrices, susceptibility distortion correction when available, and co-registrations to anatomical and template spaces). Gridded (volumetric) resamplings were performed using *antsApplyTransforms* (ANTs), configured with Lanczos interpolation to minimize the smoothing effects of other kernels (Lanczos, 1964). Non-gridded (surface) resamplings were performed using *mri_vol2surf* (FreeSurfer).

#### 2.4.2 Correlational PPI estimation

We estimated the first-level whole-brain psychophysiological interaction (cPPI) matrix in each subject’s native space. Note that, in contrast with the traditional PPI technique, cPPI results in symmetrical, *undirected* connectivity matrices. We used a region-of-interest (ROI)-based approach, obtaining connectivity terms for |N| = 416 ROIs corresponding to the cortical Schaeffer parcellation (|N| = 400) (Schaefer et al., 2018) and eight bilateral ROIs from the *aseg* atlas (accumbens, amygdala, caudate, pallidum, putamen, thalamus, hippocampus anterior and posterior) (Fischl et al., 2002). The different conditions (“tasks”) of interest were modeled based as events with onsets and durations corresponding to the experimental trial period (i.e., 2 seconds epochs that comprised the entire period of picture presentation). The task regressors were convolved with a double-gamma canonical hemodynamic response function (HRF). Events were assigned to a given condition based on the participant’s response during the subsequent memory test namely: Source (subsequent item-source association [Yes response to Q1 and Q2 and correct response to Q3]), Item (subsequent item memory without memory for the association [correct Yes response to Q1 and either a No response to Q2, or incorrect response to Q3]), Miss memory trials, and trials with no response. BOLD data (average time series) for the 416 ROIs were deconvolved into estimates of neural events (Gitelman et al., 2003). Each task time course from the first-level activity GLM design matrix was multiplied separately by the deconvolved neural estimates from the seed region and convolved with a canonical HRF, creating PPI terms. Pairwise (ROI-to-ROI) partial Pearson’s correlations were estimated for each participant separately by correlating the ROI-specific source memory PPI terms while controlling for (i) the remaining PPI regressors, (ii) the observed BOLD signal in both regions, and (iii) the original HRF-convolved task regressors. All correlation coefficients were Fisher-transformed to z values. Finally, cPPI values were demeaned within-individual to account for non-neural effects in the implicit baseline. The individual source-connectivity matrices were used for higher-level analysis. For illustrative and communication purposes, the ROIs were grouped based on 18 networks (subcortical network plus 17 cortical networks as defined by Yeo and colleagues (2011).

#### 2.4.3 Longitudinal structural preprocessing

For the structural longitudinal analysis, we performed cortical reconstruction and volumetric segmentation of the T1w scans using the longitudinal FreeSurfer stream v.6.0 (Reuter et al., 2012) (http://surfer.nmr.mgh.harvard.edu/fswiki). The images were initially processed using the cross-sectional stream thoroughly described elsewhere (Dale et al., 1999; Fischl and Dale, 2000; Fischl et al., 1999). The automatized processing pipeline includes removal of nonbrain tissues, Talairach transformation, intensity correction, tissue and volumetric segmentation, cortical surface reconstruction, and cortical parcellation. Next, an unbiased within-subject template volume based on all cross-sectional images was created for each participant, using robust, inverse consistent registration (Reuter et al., 2010). The processing of each time point was then reinitialized with common information from the within-subject template, significantly increasing reliability and statistical power. Before group analysis, cortical hemispheres were brought to *fsaverage* space and surface smoothing was applied at 12 mm FWHM. For subcortical structures (i.e., hippocampi) mean bilateral volume for specific structures was used in the analyses.

### 2.5 Higher-level analysis

#### 2.5.1 Main effects of whole-brain correlational PPI

We ran four GLM models on the whole-brain connectivity cPPI matrices to assess the mean connectivity and the effects of Age, Performance and Age×Performance interaction. Sex was used as a covariate of no-interest in all models. The models were built in a step-wise manner, adding complexity in each model. First, we assessed the mean patterns of task-dependent connectivity during memory encoding. Next, we added an age regressor to test for changes in encoding-connectivity with age. The third model tested the relation of performance (as defined by the corrected source memory scores) on encoding connectivity, age controlled. In the fourth model, we tested the Age×Performance interaction by adding the interaction regressor. This later model was restricted to edges showing a main effect of Performance. All analyses were corrected for multiple comparisons via cluster correction routines from the Network Based Statistics (NBS) toolbox (Zalesky et al., 2010), with p < 0.01 cluster-forming threshold and p < 0.025 (2 comparisons) cluster significance as determined by permutation testing (n = 5000).

#### 2.5.2 Spatial relationship between connectivity maps and term-based meta-analyses

To study the relationship between connectivity patterns and cognitive processes, we compared the topology of the main effects of Age, Performance and Age×Performance interaction with the meta-analytic patterns of activity that were associated with specific cognitive processes.

For connectivity, we estimated the “significance degree” of each ROI in the cPPI graphs; that is the number of connections (“edges”) that were significant for a given ROI in a given contrast (Age, Performance, and Age×Performance interaction). The connectivity output for each contrast was a |N| = 416 ROIs map representing the degree to which each region was related to Age, Performance, and Age×Performance interaction effects.

The meta-analytic cognitive maps were computed with the *NiMARE* package (Salo et al., 2018), which uses core functions from *Neurosynth* (Yarkoni et al., 2011). The software computes meta-analytical maps based on (mostly) activity contrasts in fMRI studies using automated text mining and coordinate extraction tools. We restricted the meta-analytical terms to those that overlapped between the Neurosynth database and the cognitive atlas (Poldrack et al., 2011) (|N| = 123 terms) thus restricting terms to specific “mental processes” (cognitive and emotional). Coordinate-based multilevel kernel density analysis (MKDA) models were used to model the specificity of the cognitive processes on neuroimaging data (Wager et al., 2009). Specificity refers to the probability of a cognitive term occurring given activation in a specific brain area. We set a term frequency threshold = 0.001 and a kernel radius = 10 mm. The remaining parameters were left to default. For comparison with “significance degree” from connectivity, the resulting meta-analytical maps were parcellated into |N| = 416 ROIs using a volumetric parcellation.

The relationship between the “significant degree” and the meta-analytical cognitive maps was assessed using Pearson’s correlations. Permutation-based significance testing (p ≤ 0.01) was performed with the *BrainSMASH* package (Brain Surrogate Maps with Autocorrelated Spatial Heterogeneity) (Burt et al., 2020). *BrainSMASH* enables statistical testing of spatially correlated brain maps by simulating surrogate brain maps with a spatial autocorrelation that matches the target map; here the meta-analytical cognitive maps (Viladomat et al., 2014). Surrogate maps (n =5000) were generated based on a Euclidean distance matrix of the center-of-gravity ROI coordinates. A null distribution was then defined by correlating the surrogate and the “significant degree” maps.

#### 2.5.3 Relationship between connectivity patterns in older-age and brain structural decline

We studied the relationship between encoding connectivity and brain atrophy and cortical thinning in a subsample of older individuals with retrospective longitudinal data (n = 81, age > 50 years). The longitudinal data spanned back on average 8.1 (SD = 0.93) years; the timing of the last observation overlapped with the timing of the encoding task. We focused on the clusters that were associated with memory performance with increasing age. We used a summarized metric that consisted of mean encoding connectivity from the clusters identified in the Age×Performance interaction models (see above for more details). Hereafter, we refer to those metrics as *memory-positive* and *memory-negative in older age*, as the resulting clusters were associated either with higher and lower memory performance with higher age, respectively. We tested whether these patterns of connectivity were associated with whole-brain cortical thinning using Spatiotemporal Linear Mixed Effect (LME) Modelling as implemented in Freesurfer (Bernal-Rusiel et al., 2013). LME models were run as a function of Time (years from the experimental task), Connectivity and the Connectivity×Time interaction. Sex, Estimated Intracranial Volume (eICV), and Baseline Age (last measurement) were introduced as covariates of no-interest and subject identifiers as random intercepts (Bernal-Rusiel et al., 2013). Statistical significance was tested at each cortical vertex and the resulting maps were corrected for multiple comparisons using False Discovery Rate (pFDR < 0.01). Finally, we investigated whether these patterns of connectivity were associated with decreased hippocampus volume, using the same model described above.

#### 2.5.4 Relationship between connectivity patterns in older age and cognitive decline

We studied the relationship between encoding connectivity and decline in memory function and general cognition in a subsample of older individuals with retrospective longitudinal cognitive data. The longitudinal data spanned back on average 7.42 years (SD = 1.94); the last observation corresponded in time with the current experimental task. We selected total learning score from the California Verbal Learning Test (CVLT), and the Vocabulary and Matrix Reasoning test from the WASI-II battery as proxies for memory function, crystalized and fluid intelligence. To explore whether connectivity in the *memory-positive* and *memory-negative in older age* clusters were associated with cognitive decline over time we ran LME analyses as detailed above with the cognitive measures fitted as a function of Time (years from the experimental task), Connectivity and the Connectivity×Time interaction (pFDR < 0.01). Sex, and Baseline Age were introduced as covariates of no-interest and subject identifiers as random intercepts.

## 3. Results

### 3.1 Behavioral results

Memory performance in the fMRI task showed a non-linear negative relationship to age, accelerating in the sixth decade of life (F = 52.26 [p < 0.001]). The different cognitive measures were related to age (all p’s < .001), controlling for sex; higher age was related to lower memory and visuospatial reasoning and higher vocabulary performance. See **Fig. 2** and **Supplementary Table 2** for additional information.

**Fig.2.**
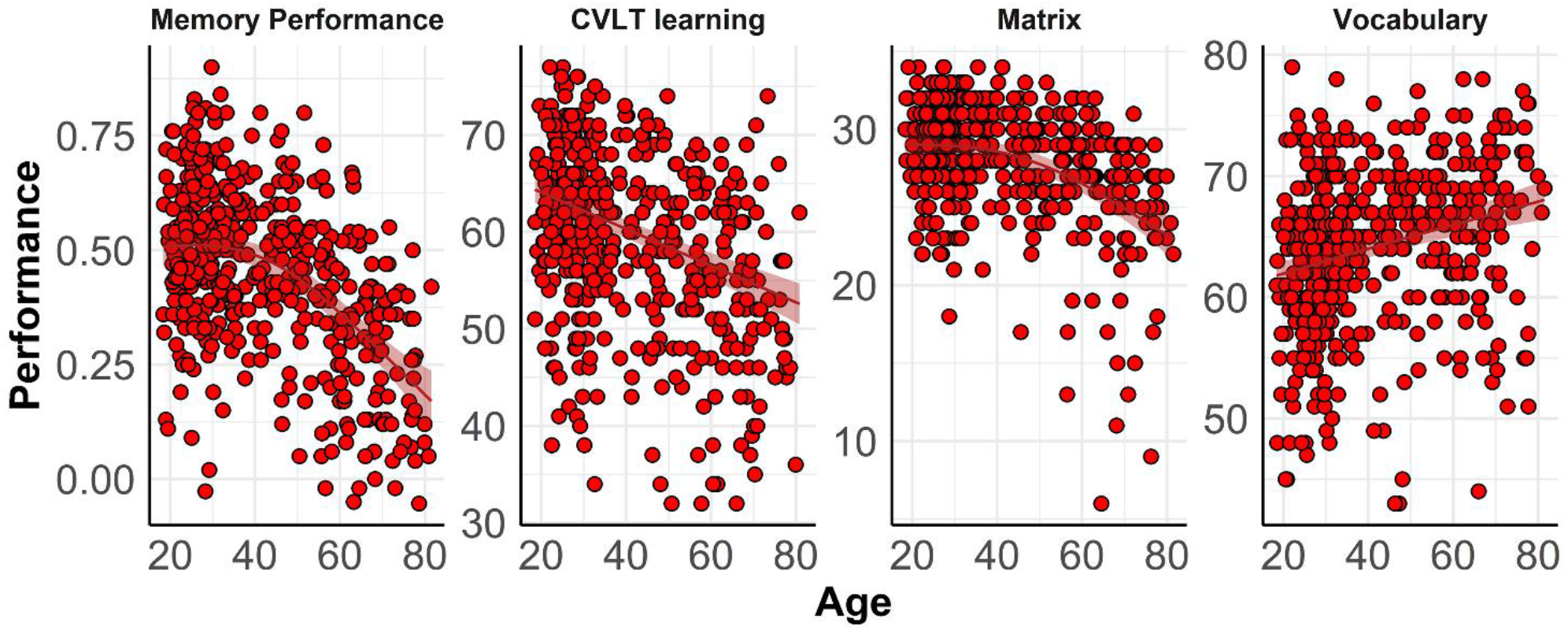
Trajectories of cognition throughout the adult lifespan. Cognitive performance was fitted by age using Generalized Additive Models (GAMs), controlled for sex. Memory performance = corrected source memory score from the experimental task; CVLT learning = words learned and recalled across the five CVLT learning trials; Vocabulary and Matrix = WAIS-IV vocabulary and matrices reasoning raw scores. Ribbons represent 95% confidence intervals.

### 3.2 Whole-brain encoding connectivity

In the main analyses, we assessed the effect of the mean task connectivity patterns during memory encoding and their association with Age, Performance (corrected source memory score from the experimental task), and the Age×Performance interaction.

#### 3.2.1 Mean encoding connectivity

Mean across-participants connectivity during encoding (**Fig. 3A**) was characterized by high intranetwork connectivity values and high internetwork connectivity between default-mode subnetworks and somatomotor networks. Low internetwork connectivity of salience, control, and limbic networks with subcortical, visual, somatomotor, and dorsal attention networks was seen.

**Fig.3.**
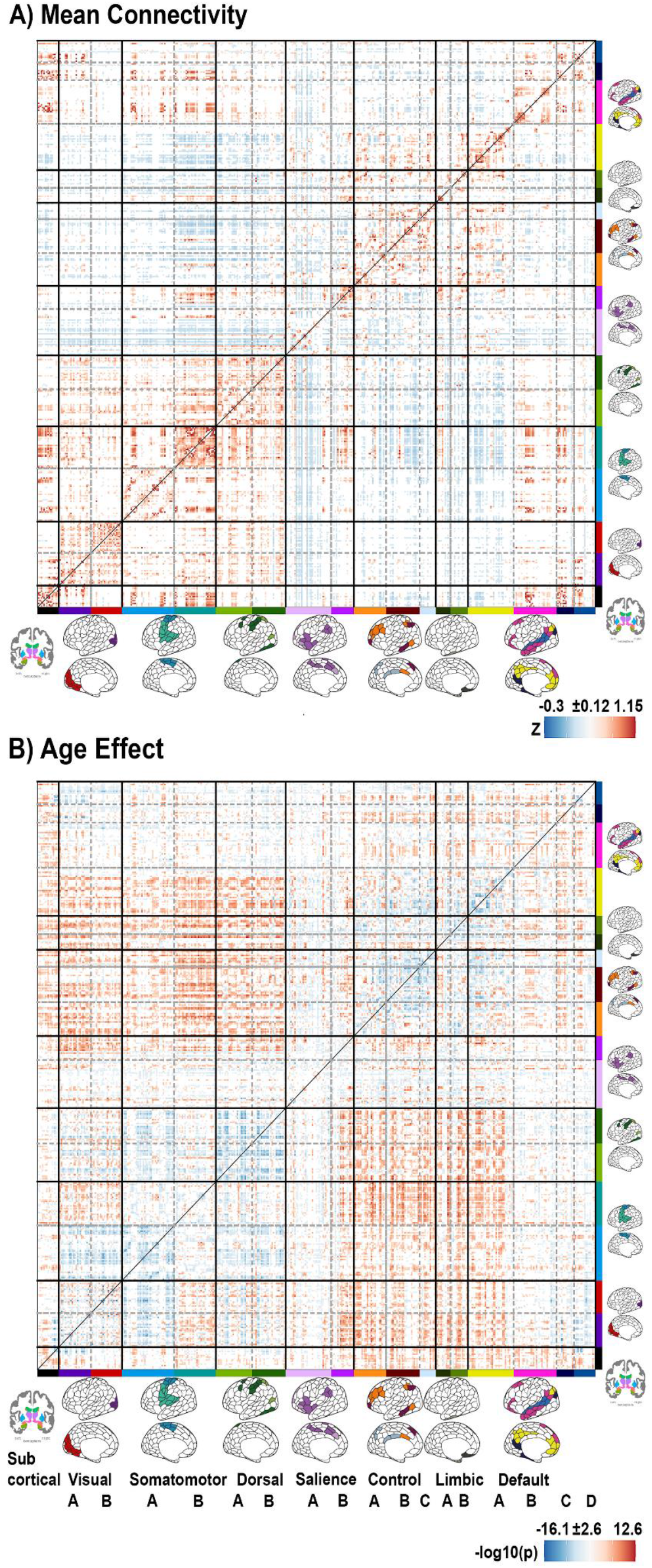
P-values matrices for mean encoding connectivity and age effects. ROIs were grouped based on the Yeo-17 atlas (Yeo et al., 2011) and a subcortical network. Default D = TemporoParietal Network. Red represents higher connectivity values and positive age effects, while blue represents lower connectivity values and negative age effects. For the mean effect, only connections ≥ 1 SD of the mean are displayed. For the age effects, only connections in FWE-corrected significant clusters are displayed (p < 0.01).

#### 3.2.2 Age effects

Pairwise connectivity between regions involved in higher cognitive functions and unimodal and attentional regions increased with higher age. Specifically, with age connectivity was higher between control, limbic, and default-mode subnetworks with somatomotor, visual, subcortical regions, and the dorsal attentional stream. Conversely, intranetwork connectivity was lower in older adults. See **Fig. 3B**. As resulted from the Mantel test, we found an inverse relationship (r = −0.19, p < 0.001 from Mantel test [n = 10000 permutations]) between the matrices of mean encoding connectivity and matrices of age effects, suggesting a likely “dedifferentiation” of the connectivity patterns with higher age. The inclusion of Performance in the model did not qualitatively affect the results.

#### 3.2.3 Performance effects

Better memory performance (positive performance) in the task was associated with higher connectivity independently of age within posterior parietal and frontal regions, namely the superior parietal lobule, auditory and somatomotor areas, the frontal operculum and medial prefrontal regions (age, sex controlled; **Fig. 4B**). Conversely, poorer memory performance (negative performance) was associated with higher connectivity within posterior lateral default-mode network regions, medial default-mode network regions, dorsal prefrontal areas, lateral prefrontal areas, and the temporal pole (**Fig. 4C)**.

**Fig.4.**
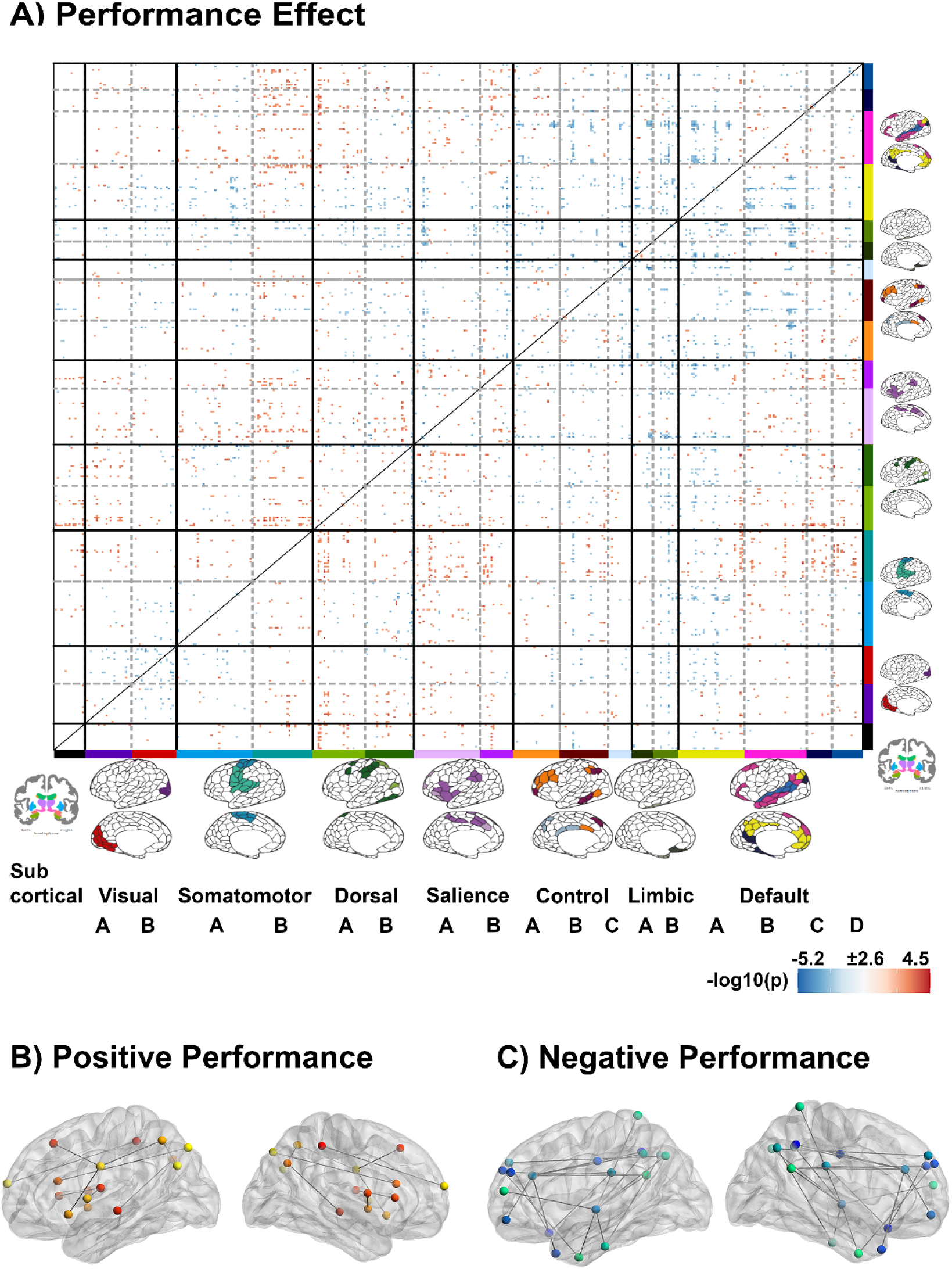
Performance effects. **A** P-values matrix for performance effects. Performance was defined as (corrected) source memory performance in the experimental task. ROIs were grouped based on the Yeo-17 atlas (Yeo et al., 2011) and a subcortical network. Default D = TemporoParietal Network. Red represents higher connectivity values and positive performance effects and vice versa for the blue scale. For the performance effects, only connections within FWE-corrected significant clusters are displayed (p < 0.01). **B-C** Top 5% of significant nodes shown overlaid to 3D-brain BrainNet Viewer (Xia et al., 2013 http://www.nitrc.org/projects/bnv/). Nodes are filled with red to yellow scales from lower to higher connections (and blue to turquoise from lower to higher connections) that indicate the number of connections (“significance degree”). Only edges between drawn nodes are displayed.

#### 3.2.4 Age×Performance interaction effects

Next, we tested whether there were Age×Performance interaction effects within the regions showing a main effect of performance. We found two significant (FWE controlled) clusters showing positive and negative Age×Performance interactions, respectively. See **Fig. 5** for a visual illustration. The first cluster (*memory-positive in older age*) included connectivity between medial temporal and posterior parietal regions, including the retrosplenial cortex, the inferior and superior parietal lobules, and regions in the medial temporal lobe. Higher connectivity between these regions was associated with better performance with higher age (**Fig. 5B**). The second cluster (*memory-negative in older age*) corresponded to connections between frontal, parietal, and visual regions. Increased connectivity between these regions was associated with lower performance in older participants (**Fig. 5C**).

**Fig.5.**
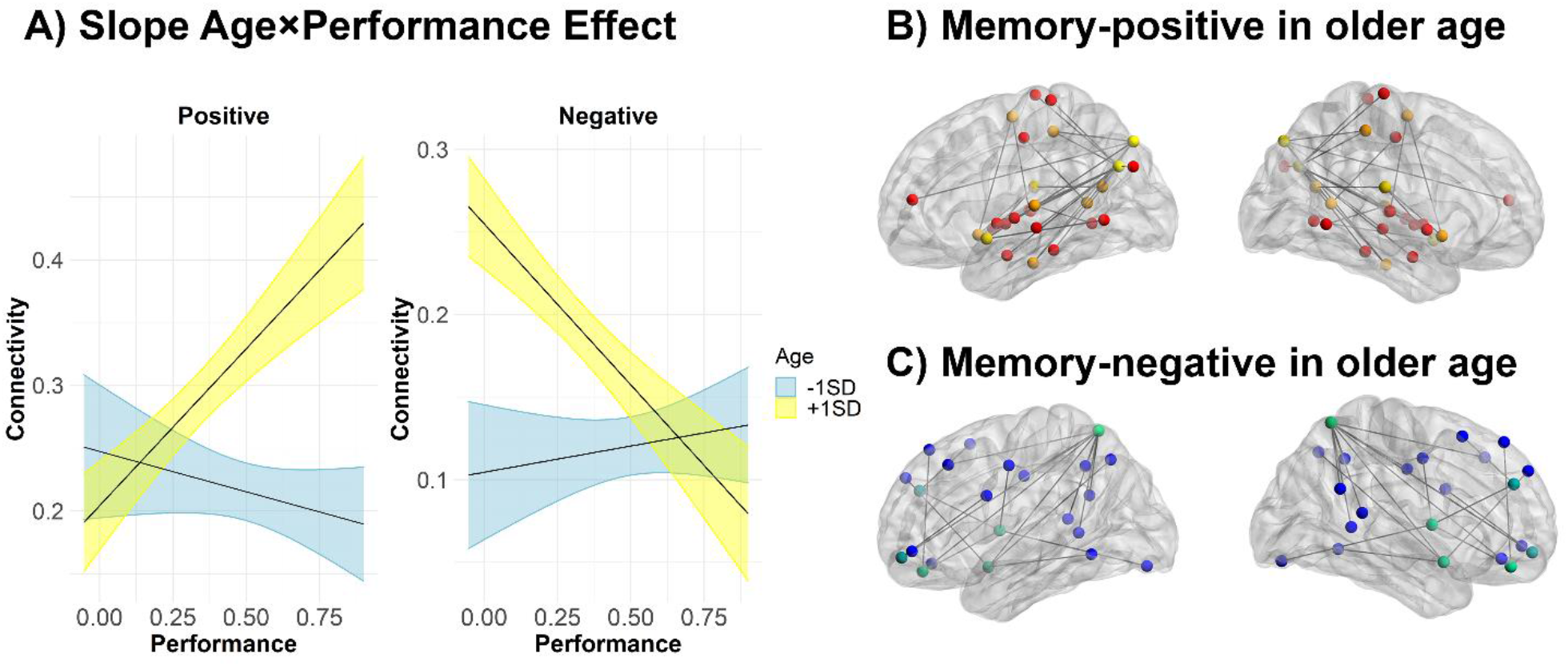
Age×Performance interaction effects. **A** Relationship between performance and task-dependent connectivity across age. For illustrative purpose, the effects of performance were predicted at two levels (± 1 SD Age; mean Age = 41.65 [SD = 17.18] years). Note though that Age was introduced as a continuous regressor in the model. Ribbons represent 95% confidence intervals. **B-C** Top 5% of significant nodes shown overlaid to 3D-brain BrainNet Viewer (Xia et al., 2013 http://www.nitrc.org/projects/bnv/). Nodes are filled with red to yellow scales from lower to higher connections (and blue to turquoise from lower to higher connections) that indicate the number of connections (“significance degree”). Only edges between drawn nodes are displayed.

### 3.3 Spatial relationship between connectivity maps and term-based meta-analyses

Next, we tested the spatial relationship between the encoding connectivity patterns and the meta-analytic activity maps associated with specific cognitive processes (p ≤ 0.01 using a permutation-based approach). We used a “significance degree” (*number of significant connections for a given ROI in a given contrast*) and specificity metrics for connectivity patterns and cognitive maps, respectively. This was done for all connectivity effects of interest (Age, Performance, and Age×Performance interaction). See **Fig. 6** and **Supplementary Table 3** for the full results. The connectivity patterns where greater connectivity was related to higher age (older age) overlapped significantly with the patterns characterizing retrieval, recall, and encoding processes. Conversely, the connectivity patterns where greater connectivity was associated with younger age overlapped with maps related to imagery, spatial attention, and movement activity. The connectivity patterns where greater connectivity was associated with better memory performance overlapped with multisensory, integration and speech production areas. Connectivity patterns where greater connectivity was related to worse performance overlapped with maps associated with salience, emotion, and belief. The spatial patterns of connectivity associated with positive Age×Performance interaction (*memory-positive in older-age*) overlapped with the activity patterns associated with mental imagery. No terms were associated with negative Age×Performance interaction (*memory-negative in older age*). These results inform us on cognitive processes that may be related to successful memory performance in successful aging, that is, integrative and multisensory strategies and mental imagery.

**Fig.6.**
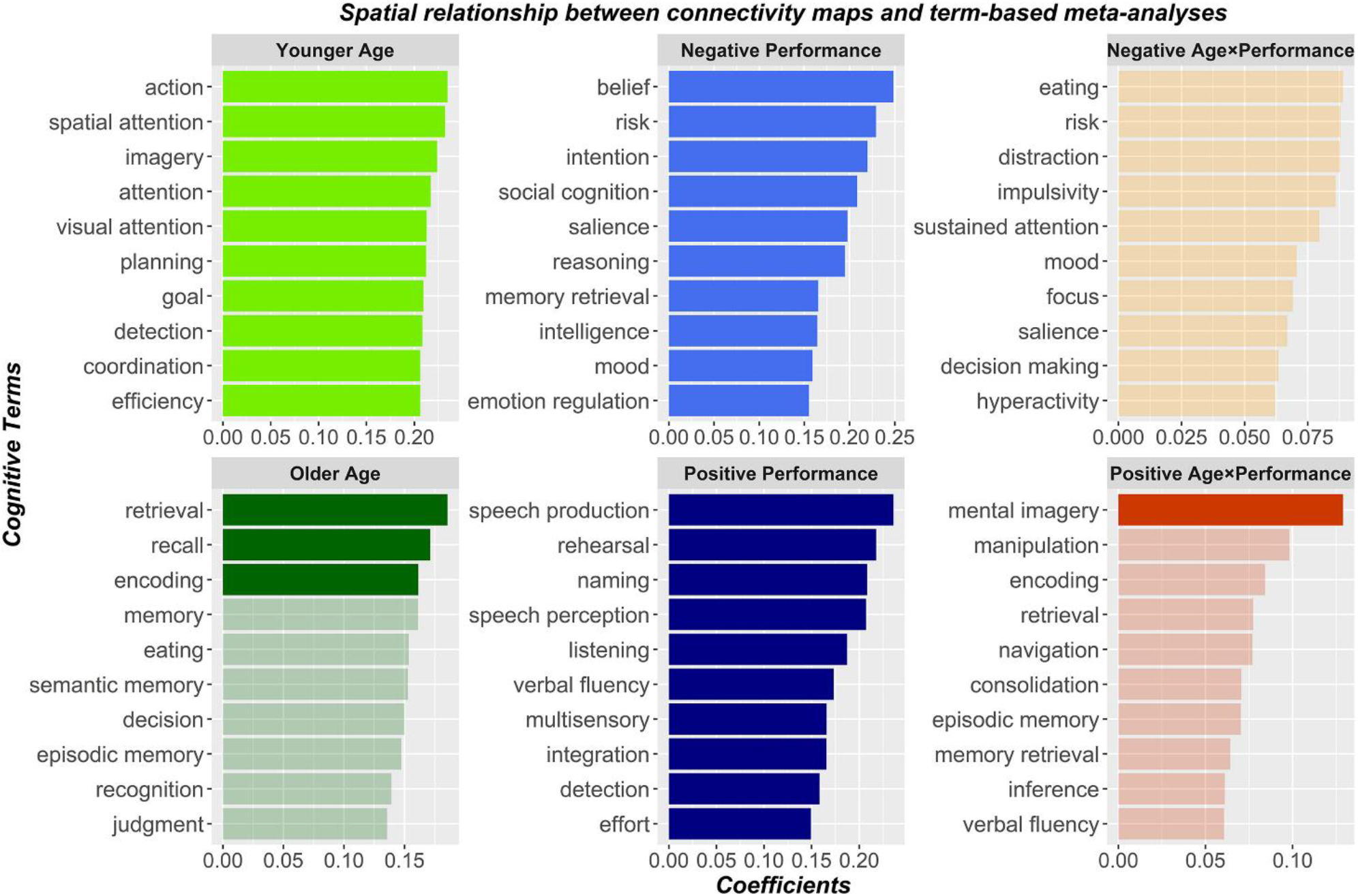
Spatial relationship between connectivity maps and meta-analytic patterns associated with specific cognitive processes. We displayed the top cognitive terms associated with each contrast. Opaque colors reflects terms that survived the significance threshold (p ≤ 0.01) as determined by a permutation approach using BrainSMASH (Burt et al., 2020). X-axis represents the empirical Pearson’s correlation (r), note that different ranges are depicted for each contrast.

### 3.4 Relationship between connectivity patterns in older-age and brain structural decline

We assessed the relationship between brain atrophy and cortical thinning and connectivity patterns in older age to investigate whether age-related connectivity changes reflected maintenance or compensatory responses. This analysis was performed in a subsample of older participants (n = 81, age >50) with retrospective longitudinal neuroimaging data (see **Supplementary Table 1**). A Linear Mixed Effects analysis (controlled for Sex, eICV, and Baseline Age) revealed that the cluster related to higher memory performance in older age (*memory-positive in older age*) was associated with less hippocampal volume decrease over time (F = 44.2, p < 0.001; **Fig.7A**). In contrast, the cluster associated with lower memory performance in older age (*memory-negative in older age*)was related to a steeper volumetric decline of hippocampi (F = −65.71, p < 0.001; **Fig.7B**) and cortical thickness decline over time. Indeed, the cortical analysis showed a positive association between connectivity in this *memory-negative in older age* cluster and cortical thinning in two small clusters encompassing (1) the left precentral and (2) the anterior fusiform and the entorhinal cortices (pFDR < 0.01, **Fig.7C**). Overall, the results supported the hypothesis that the patterns of connectivity associated with higher and lower performance in older age were related to structural maintenance versus decline of brain regions involved in memory processes.

**Fig.7.**
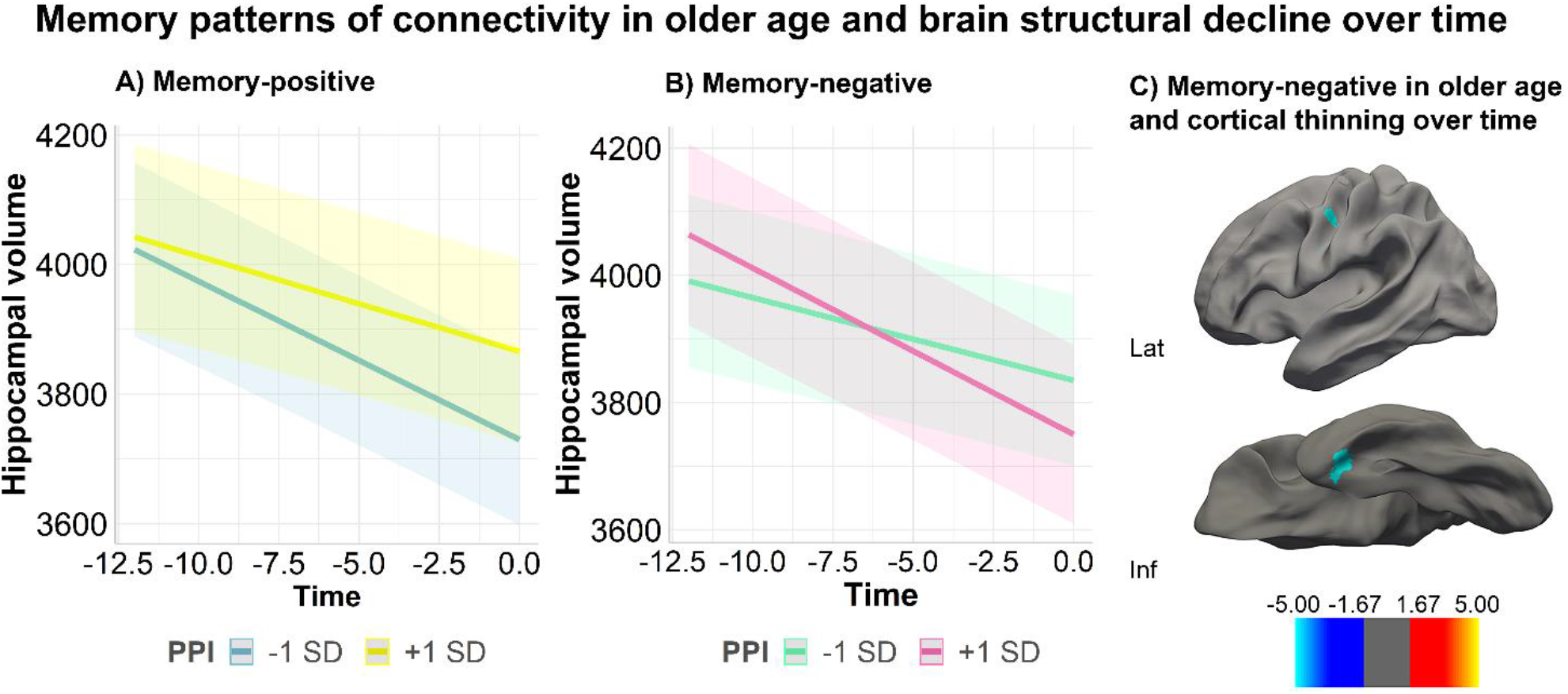
Relationship between memory pattern of connectivity in older age and decline of brain structures over time. A-B Relationship between memory patterns of connectivity in older and longitudinal structural hippocampal volume. **A** The yellow line represents higher memory performance in memory-positive cluster. **B** The pink line represents lower memory performance in memory-negative cluster. Results are significant at pFDR < 0.01. **C** Whole-brain cortical thickness decline and memory-negative in older age. Only regions showing significant thinning over time are shown. Left hemisphere. Maps are corrected for pFDR < 0.01. In the colorbar -log10(p) values are displayed, red represents higher values, blue lower values.

### 3.5 Relationship between connectivity patterns in older age and cognitive decline

We assessed the relationship between memory patterns of connectivity in older age and cognitive decline using retrospective cognitive data (age > 50). See **Supplementary Table 1** for details. LME models (controlled for Sex and Baseline Age) revealed a relationship between connectivity patterns associated with higher memory performance in older age (*memory-positive in older age*) and less decline in CVLT learning scores. However, the association did not survive multiple comparison corrections (pFDR = 0.07).

## 4. Discussion

We estimated whole-brain functional connectivity during episodic memory encoding, specifically focusing on age-related differences in connectivity and how they were associated with memory performance across the lifespan. In higher age, we found lower intranetwork and higher internetwork connectivity between regions involved in higher cognitive functions and the dorsal attention stream, sensorimotor and subcortical regions during encoding. Successful memory performance in higher age overlapped with networks involved in mental imagery. Among older adults, greater hippocampal and cortical atrophy was related to less favorable connectivity changes, reflecting maintenance processes over time.

The age effects on encoding connectivity are partially in agreement with previous rs-fMRI and task-fMRI studies, suggesting that some of the functional age-differences are task-independent. For example, several resting-state studies have found reduced intranetwork and increased internetwork connectivity, indicating that brain networks become less specialized during aging (Betzel et al., 2014; Geerligs et al., 2015). Moreover, we found higher age-related connectivity between control and dorsal attention networks, which has been reported previously in a fMRI study using a different task (Grady et al., 2016). This might be interpreted as an over-recruitment of cognitive control processes due to the cognitive demands of the task. Likewise, higher connectivity between inversely engaged networks, such as the control and the dorsal attention respectively with the default-mode, has been described in several tasks in aging (Spreng et al., 2016; Spreng and Turner, 2019). These patterns may reflect age-related features during memory tasks such as lower flexibility in shifting from external and internal attention and semantization of cognition as older individuals might rely more on acquired knowledge. Although speculative, this interpretation receives further support from our results as the connectivity patterns associated with older age mapped unto cognitive processes such as retrieval and recall, suggesting that older participants might rely more on already acquired knowledge and schematic information during the encoding task. However, our results also reveal task-specific changes in connectivity. Compared to fMRI acquired during resting-state and other cognitive domains, we also identified the spatial correspondence between the connectivity patterns and cognitive maps, informing us on the specific cognitive processes possibly involved in the task. Younger people exhibited higher connectivity between areas overlapping with regions known to support cognitive processes relevant for our encoding task such as visual attention, action, and imagery. Some of these strategies may also have been adopted by older participants that performed better, which may be interpreted to be in accordance with the maintenance process framework.

We found that the relationship between connectivity and memory performance differed as a function of age. Older people who performed better showed higher connectivity between medial temporal and posterior parietal regions, including the retrosplenial cortex. As shown in a previous study of older participants (Kaboodvand et al., 2018), episodic memory performance was positively associated with functional connectivity between the retrosplenial cortex and the medial temporal lobe. The retrosplenial cortex is a key mediator in facilitating the communication between medial temporal and other default mode networks regions, leading to memory performance success (Kaboodvand et al., 2018). In our study, the connectivity changes that were related to better performance in older participants overlapped spatially with the maps associated with mental imagery, in which the engagement of the retrosplenial cortex is widely described (Chrastil, 2018). These strategies and functional connectivity changes mimicked those associated with younger age.

The patterns of connectivity associated with successful performance in older participants were related to volumetric maintenance of the hippocampus, critically involved in memory encoding. This is in agreement with the maintenance theory of cognitive aging (Nyberg et al., 2012), as hippocampal decline is a main factor behind memory decline in older age (Gorbach et al., 2017). Conversely, we found no evidence that the pattern of connectivity associated with higher performance in older age reflected a compensatory attempt to overcome structural decline. The widespread over-recruitment of different regions in frontal, parietal, and visual areas was not related to memory performance.

Indeed, these connectivity changes were associated with lower performance and structural loss in the medial temporal lobe over time. Cognitive decline was associated with structural decline and a maladaptive organization in the functional architecture, whereas successful memory performance in older participants reflected relative structural integrity over time and functional connectivity changes that supported the use of “younger” cognitive strategies such as mental imagery.

### 4.1 Limitations and methodological remarks

fMRI during task performance is suited for investigating brain dynamics associated with specific cognitive processes as they possess experimental control while participants perform a task-of-interest (Campbell and Schacter, 2017). Despite commonalities across states, the functional brain architecture differs across task contexts (Davis et al., 2017) as brain regions reconfigure their connectivity patterns in a flexible way based on the current demands of the task (Cole et al., 2013). This point is supported in our study, as the connectivity patterns associated with successful memory performance mapped on networks involved in cognitive processes relevant for this encoding task. The main disadvantage of task-fMRI is that differences in the experimental design may hamper generalization (Damoiseaux and Huijbers, 2017). Some of our findings agree with previous rs-fMRI and task-fMRI research and thus likely represent task-invariant features of the aging brain. However, other findings, as highlighted by the spatial correlation between functional connectivity and mental imagery processes, seem more constrained to the specific demands of the task, although they relate to real-life function.

The cPPI framework allowed for a whole-brain *undirected* (symmetric) assessment of task connectivity. cPPI does not imply inferences of directionality. The cPPI connectivity values reflect correlations between regions during selected task-periods of an fMRI run, controlling for stimulus-driven co-fluctuations and intrinsic functional connectivity between the ROIs. When cPPI connectivity values estimated from different task-periods are subtracted, the resulting metric is largely comparable to traditional regression-based PPI approaches. However, when conditions are not subtracted - as in the current paper - cPPI is akin to “residualised” task-connectivity and beta series correlation, i.e. the similarity between two regions’ trial-to-trial fluctuations in BOLD amplitude during task (Di et al., 2020). The different implications of the metrics are largely omitted in the literature but have consequential implications for the interpretation, that is in this case, a task-state of integrated connectivity, instead of a shift in connectivity driven by the specific task.

The effects of demeaning data within participants are also consequential for the interpretation of our results. This approach minimizes the possibility of non-neural confounds that affect the implicit baseline being the main drivers of connectivity differences across individuals. Some of these confounds are known to be greatly correlated with age (Campbell and Schacter, 2017). However, data demeaning only allowed us to interpret the results in relative terms, and in terms of reorganization. Note that many graph-theoretical studies use thresholded, binarized data and thus face a similar problem. It is however possible that some findings are a side-effect of this step. For example, the patterns of connectivity associated with lower performance in older participants are spatially unstructured and thus might represent unspecific changes in the functional connectome rather than reduced functional connectivity amongst specific regions.

Despite the structural and cognitive retrospective longitudinal data available, the main limitation of this study is the lack of longitudinal task-fMRI, which would have allowed us to assess how the functional architecture of the brain during memory tasks changes over time.

## 5. Conclusions

This study provides novel insights in whole-brain connectivity during encoding and its relation with age, cognitive processes and structural decline in older age using a large sample encompassing the entire adulthood. Connectivity patterns underlying successful memory function in older age spatially mapped onto mental imagery processes and were related to structural brain maintenance over time. These results provide a bridge between the cognitive processes and the biological mechanisms that support memory function maintenance and decline in older age.

## Acknowledgements

This work was supported by the Department of Psychology, University of Oslo (to K.B.W., A.M.F.), the Norwegian Research Council (to K.B.W., A.M.F.) and the project has received funding from the European Research Council’s Starting Grant scheme under grant agreements 283634, 725025 (to A.M.F.) and 313440 (to K.B.W.).

