## Supplementary Tables for "Whole-brain connectivity during encoding: age-related differences and associations with cognitive and brain structural decline"

**Supplementary Table 1**
Longitudinal MRI variables

| | Tp1 | Tp2 | Tp3 | Tp5 |
|---|---|---|---|---|
| **n** | 81 | 81 | 80 | 1 |
| **Age** | 57.44 (7.89) | 61.17 (7.52) | 65.38 (7.05) | 78.63 |
| **Time** (Age from baseline *years*) | -8.01 (0.93) | -4.37 (0.49) | 0 | 0 |
| **Hippocampal volume** (*mm*$^3$) | 3957.86 (427.78) | 3886.03 (437.73) | 3795.83 (440.94) | 3437.65 |

Longitudinal cognitive variables

| | Tp1 | Tp2 | Tp3 | Tp4 | Tp5 | Tp6 |
|---|---|---|---|---|---|---|
| **n** | 94 | 94 | 94 | 11 | 11 | 6 |
| **Age** | 59.50 (9.00) | 62.75(8.06) | 66.63 (7.24) | 73.30 (2.95) | 74.23 (3.21) | 75.02 (2.96) |
| **Time** (Age from baseline *years*) | -7.42 (1.93) | -4.17 (0.71) | -0.29 (0.85) | -2.28 (0.84) | -1.35 (1.39) | 0 |
| **CVLT Total** | 53.28 (9.81) | 56.54(10.59) | 54.59 (9.31) | 60 (15.77) | 61.73 (8.56) | 61.5 (10.84) |
| **Matrix** | 26.51 (4.64) | 26.88 (5.47) | 25.82 (4.75) | / | 25.6 (0.89) | 27.5 (2.35) |
| **Vocabulary** | 65.50 (6.41) | 65.40 (6.74) | 65.99 (6.5) | / | 66.8 (6.61) | 66 (8.92) |

Descriptive statistics represent mean (SD) for the different timepoints (Tp). CVLT learning = words learned and recalled across the five CVLT learning trials; Vocabulary and Matrix = WAIS-IV vocabulary and matrices reasoning raw scores.

**Supplementary Table 2**
Main demographic, neuropsychological and behavioral variables

| **Sample descriptives** | **Mean (SD)** | **Age (F [p])** |
|---|---|---|
| Sex (female:male) | 336:152 | - |
| Age | 41.65 (17.20) | - |
| Age range | 18:81 | - |
| Memory Performance | 0.45 (0.18) | 52.26 (<0.001)* |
| CVLT learning | 58.58 (9.34) | 63.45 (<0.001)* |
| Matrix | 28 (3.75) | 35.31 (<0.001)* |
| Vocabulary | 64.07 (6.68) | 28.28 (<0.001)* |
| Item memory | 0.18 (0.08) | 6.34 (0.001)* |
| Miss | 0.22 (0.11) | 9.08 (0.002)* |
| Correct Rejections | 0.92 (0.08) | 27.59 (<0.001)* |
| False Alarms | 0.056 (0.05) | 20.98 (<0.001)* |

Descriptive statistics represent mean (SD) for quantitative variables and numbers by condition for the qualitative variables. Effects of age on each variable fitted by GAM models (sex controlled). Memory performance was defined as corrected source memory in the experimental task. CVLT learning = words learned and recalled across the five CVLT learning trials; Vocabulary and Matrix = WAIS-IV vocabulary and matrices reasoning raw scores. Item memory = “Yes” response to Question 1 and either “No” to Question 2 or incorrect answer to Question 3; miss = incorrect answer to Question 1. New items were classified either as correct rejections or false alarms.

**Supplementary Table 3**
Spatial relationship between connectivity maps and meta-analytic patterns associated with specific cognitive processes

| | Terms meta-analyses | p-value | Pearson’s r |
|---|---|---|---|
| **Negative Age** | action | 0.001 | 0.234 |
| | attention | 0.004 | 0.217 |
| | coordination | 0.008 | 0.206 |
| | detection | 0.007 | 0.208 |
| | efficiency | 0.007 | 0.206 |
| | goal | 0.007 | 0.209 |
| | imagery | 0.005 | 0.223 |
| | planning | 0.006 | 0.212 |
| | spatial attention | 0.002 | 0.232 |
| | visual attention | 0.004 | 0.212 |
| **Negative Performance** | autobiographical memory | 0.008 | 0.144 |
| | belief | <0.001 | 0.248 |
| | emotion regulation | 0.003 | 0.155 |
| | intelligence | 0.002 | 0.164 |
| | intention | <0.001 | 0.219 |
| | knowledge | 0.007 | 0.144 |
| | memory retrieval | 0.002 | 0.165 |
| | mood | 0.003 | 0.159 |
| | psychosis | 0.007 | 0.142 |
| | reasoning | <0.001 | 0.194 |
| | risk | <0.001 | 0.229 |
| | salience | <0.001 | 0.197 |
| | semantic memory | 0.008 | 0.139 |
| | social cognition | <0.001 | 0.208 |
| | thought | 0.004 | 0.155 |
| **Positive Age** | encoding | 0.009 | 0.161 |
| | recall | 0.006 | 0.171 |
| | retrieval | 0.002 | 0.185 |
| **Positive Interaction** | mental imagery | 0.009 | 0.129 |
| **Positive Performance** | competition | 0.008 | 0.151 |
| | detection | 0.007 | 0.158 |
| | effort | 0.007 | 0.149 |
| | integration | 0.007 | 0.165 |
| | listening | 0.001 | 0.187 |
| | multisensory | 0.003 | 0.165 |
| | naming | <0.001 | 0.208 |
| | reading | 0.010 | 0.144 |
| | rehearsal | <0.001 | 0.217 |
| | speech perception | <0.001 | 0.207 |
| | speech production | <0.001 | 0.236 |
| | verbal fluency | 0.005 | 0.173 |
| | word recognition | 0.008 | 0.148 |

Spatial relationship between connectivity maps and meta-analytic patterns associated with specific cognitive processes**.** See Fig.6 in the text for a visual description. We displayed, in alphabetic order, the top cognitive terms associated with each contrast, that survived the significance threshold ($p \leq 0.01$) as determined by a permutation approach using BrainSMASH (Burt et al., 2020). Pearson’s correlation coefficients are displayed.
